# Deep-intronic variants in *USH2A*: the importance of model selection in variant interpretation and pathogenicity assessment

**DOI:** 10.64898/2026.10.28.753787

**Authors:** Kim Rodenburg, Renske T.W. Schellens, Sanne Broekman, Leony Fenwick, Florian A. Winkelaar, Dirk H. Wijn, Galuh D.N. Astuti, Gelana Khazeeva, Christian Gillissen, Frans P. M. Cremers, Ronald J.E. Pennings, Theo Peters, Ulrich Zechner, David Rosenkranz, Christian Betz, Monika Gawai, Vanessa Hochleimer, Suzanne E. de Bruijn, Erik de Vrieze, Hanno J. Bolz, Susanne Roosing, Erwin van Wijk

## Abstract

Interpreting splice-altering *USH2A* variants is challenging when these result in the alteration of splicing in only a fraction of transcripts. We investigated two deep-intronic *USH2A* variants from a single case, the inherited variant c.5573-834A>G and the *de novo* variant c.11047+345A>G, using a minigene splice assay and transcript analysis in patient-derived photoreceptor precursor cells and nasal epithelial cells. The minigene assay revealed the inclusion of a pseudoexon in the presence of the c.11047+345A>G variant, but did not reproduce previously reported aberrant splicing for c.5573-834A>G. In patient-derived cells, c.5573-834A>G resulted in the inclusion of a pseudoexon in part of the transcripts, whereas c.11047+345A>G did not result in detectable levels of aberrantly spliced transcripts.

To assess the impact of residual wildtype transcripts on retinal function, we used the previously developed humanized *ush2a^PE40+M^*^1^ knock-in zebrafish model which allows the modeling of the impact of mixed wildtype and pseudoexon-containing *ush2a* transcripts. This model demonstrated that detectable levels of usherin expression are maintained even when only a fraction of wildtype transcripts lacking pseudoexon inclusion remain. Collectively, these findings advocate for a more nuanced classification of partially splice-altering *USH2A* variants. The mere presence of alternative transcripts should not be considered sufficient evidence for pathogenicity. Within this framework, data on the c.5573-834A>G variant do not support a (likely) pathogenic classification, and findings related to the c.11047+345A>G variant are consistent with a likely benign classification.

## Introduction

Usher syndrome (USH) is the leading cause of hereditary deaf-blindness in humans and has a prevalence of 1:15,000 individuals worldwide. USH presents with an autosomal recessive mode of inheritance and can be distinguished in four distinct clinical types (USH1-4) based on the age at which the different clinical symptoms become apparent, the severity and rate of progression of the symptoms, and the presence or absence of vestibular dysfunction. Of the clinical types, USH type 2 is the most prevalent and is typically characterized by a moderate to severe bilateral sensorineural hearing impairment from birth and bilateral progressive loss of visual function due to retinitis pigmentosa, with an onset typically in the second decade of life or later^1–3^. Of all USH type 2 cases, 55-90% is explained by pathogenic variants in *USH2A*^4,5^.

Over 1,760 unique (likely) pathogenic *USH2A* variants have been reported to date (Leiden Open Variation Database V.3.0, accessed August 2026), ranging from single-nucleotide variants (SNV), such as missense and nonsense variants, to structural variants (SVs). Following the implementation of short-read whole genome sequencing (WGS), an increasing number of deep-intronic variants affecting pre-mRNA splicing have been identified that are associated with USH or non-syndromic retinitis pigmentosa^6–8^.

The pathogenic c.7595-2144A>G variant was the first deep-intronic *USH2A* variant identified to generate a strong splice donor site and results in the exonification of 152 nucleotides of intron 40 (PE40)^9^. In patient-derived cells, this variant results in PE40 inclusion in 100% of *USH2A* transcripts, resulting in the premature termination of usherin translation (p.Lys2532Thrfs*56)^10^. In contrast, other deep-intronic *USH2A* variants have been reported with a less profound effect on pre-mRNA splicing, resulting in the inclusion of pseudoexons in a portion of transcripts while also retaining wildtype transcripts^11^. The clinical relevance and the impact these variants have on protein levels remains unclear, and these variants are therefore generally classified as variants of unknown significance (VUS).

A widely used method to assess the impact of splice-altering variants is the use of an artificial *in vitro* minigene splice assay. This method provides a rapid, non-invasive and cost-effective strategy to assess the impact of a variant on pre-mRNA splicing in a non-native and non-personalized context as an alternative approach to this strategy, photoreceptor precursor cells (PPCs) generated from patient-derived induced pluripotent stem cells (iPSCs) can be used to assess the effect of variants associated with inherited retinal diseases such as USH on splice alteration^11,12^. PPCs enable the evaluation of a genetic variant in the entire genetic context of the affected individual in cells that are closest to the cells of the affected tissue and that endogenously express the gene of interest. As relevant tissue-specific splice factors are present, this approach provides a context that is biologically more relevant compared to *in vitro* minigene splice assays for observing the impact of the selected variants.

Alternatively, to evaluate the functional impact of an *in vivo* variant, a zebrafish model can be used. The zebrafish has emerged as a widely accepted translational model to study *USH2A*-associated RP^13–15^. The translational potential of this model was exemplified in a proof-of-concept study in which antisense oligonucleotides (QR-421a/ultevursen) induced the targeted in-frame skipping of the frequently mutated *ush2a* exon13 resulting in a partial restoration of usherin protein production, sufficient for normal visual function in this model^12^.

In the current study, we compared the effect of two novel deep-intronic *USH2A* variants on pre-mRNA splicing using in vitro minigene splice assays, patient-derived PPCs as well as patient-derived nasal epithelial cells. As the pathogenicity of both variants remained uncertain, we employed zebrafish as a functional model to establish a minimum threshold level of residual wildtype *USH2A* transcripts required for normal retinal function. The results of this study will aid in *USH2A* deep-intronic variant interpretation and classification.

## Methods and materials

### Individual in this study

The study adhered to the tenets of the Declaration of Helsinki and was approved by the local ethics committees of the Radboud University Medical Center (Nijmegen, The Netherlands) and the University Hospital of Cologne. Written informed consent was obtained from the participating individual prior to inclusion in this study.

### Short-read whole genome sequencing

Genomic DNA from the proband was isolated from peripheral blood lymphocytes according to standard procedures. Short-read WGS was performed at BGI Tech Solutions (Warszawa, Poland) on a BGISeq500 using 2x 150 base pair (bp) paired-end reads with a minimal median coverage of 30-fold per genome. Sequencing reads were mapped to the Human Reference Genome build GRCh38/hg38 using Burrows-Wheeler Aligner V.0.7814^16^. SNVs and small indels (<50 bp) were called using Genome Analysis Toolkit HaplotypeCaller (Broad Institute), SVs were called using Manta Structural variant Caller, based on read-pair evidence and read-depth evidence^17,18^. Copy number variants (CNVs) were called using Canvas Copy Number Variant caller, based on read-depth evidence^19^. SNVs, small indels, SVs and CNVs were annotated using an in-house pipeline as described previously^20^.

### Variant prioritization and analysis

WGS data of the proband was analyzed to identify potential pathogenic variants. Coding and non-coding SNV variants were selected based on a minor allele frequency of <0.01 in the GnomAD population database (all populations) (V3.1.2, with manual checks in V4.1.1 for selected variants)^21^. The selected rare variants were prioritized based on protein effect. Nonsense, start- or stop-affecting variants, as well as frameshift, in-frame, missense variants, canonical and intronic splice variants were further examined in more detail. For missense variants, CADD_PHRED and REVEL scores were obtained and variants meeting the predefined thresholds of both in silico prediction tools (CADD_PHRED: >15, range 0-99; REVEL: >0.3, range 0-1) were selected for further investigation as possibly pathogenic candidates^22,23^. Variants meeting the threshold of a single tool were retained as lower-priority candidate variants. SpliceAI delta scores (https://spliceailookup.broadinstitute.org/) were obtained for all variants with a potential effect on splicing (canonical splice variants, missense, synonymous and intronic variants). Variants were considered for further testing if one out of the four provided scores (acceptor gain (DS_AG), acceptor loss (DS_AL), donor gain (DS_DG) or donor loss (DS_DL)) provided a delta score of > 0.2 using the following window setting: upstream and downstream 500 bp)^24^. Alamut™ Visual plus (V2.0) was used to visually inspect the genomic context of the positions predicted by SpliceAI delta scores. Coding SV and CNV data was prioritized on a minor allele frequency of <1% in the 1,000 genome database^25^. Inversions and duplications were only considered potentially pathogenic when disrupting an RP- or USH-associated gene as proposed by De Bruijn et al.^26^.

### Long-read whole-genome sequencing

Short DNA fragments were removed by precipitation (SRE kit, Pacific Biosciences, Menlo Park, CA), followed by shearing to approximately 16 kb DNA size fragments by robot liquid handling (Hamilton NGS Star, Hamilton, Bonaduz CH). Whole-genome library preparation and polymerase binding were done following the SMRTbell prep kit 3.0 protocol (Pacific Biosciences, Menlo Park, CA). Sequencing was performed on a PacBio Revio® system with one sample per SMRT cell, 260 pM loading, 24h run time. Typical outputs exceeded 80 Gb of HiFi reads. Secondary analysis for SNV, CNVs and SVs was conducted with an in-house pipeline integrating available PacBio tools developed for HiFi long-read analyses, providing information also on phasing, methylation status, and repeat expansions, as reported previously^27^.

### Minigene splice vector

Minigene constructs were generated as previously described^28^. In short, the genomic regions containing the *USH2A* variants c.5573-834A>G and c.11047+345A>G, along with at least 450 base pairs flanking each variant position, were amplified from the proband’s genomic DNA and cloned into pDONR201 (Invitrogen, Waltham, MA, USA). The complete pDONR201 constructs were sequence verified using Sanger sequencing and subsequently cloned into destination pCl-NEO vector (Thermo Fisher Scientific, Carlsbad, CA, USA) containing human *RHO* exon 3 and exon 5 using Gateway cloning technology. The resulting individual four minigene splicing vectors contain either *USH2A* wildtype genomic inserts or inserts harboring variants c.5573-834A>G or c.11047+345A>G. Sequences of all primers used are listed in Supplementary table 1.

### Cell culture and transfection of *in vitro* minigene splice vectors

HEK293T cells were cultured in Dulbecco’s modified Eagle’s medium (DMEM) (#D0819, Sigma-Aldrich, St. Louis, MO, USA) supplemented with 10% (v/v) fetal bovine serum (#F7524,Sigma-Aldrich, St. Louis, MO, USA), 1% penicillin-streptomycin (#P4333, Sigma- Aldrich, St. Louis, MO, USA) and 1% sodium pyruvate (#S8636, Sigma-Aldrich, St. Louis, MO, USA). Cells were passaged twice per week upon standard trypsinization (#DF0152-15-9, Thermo Fisher Scientific, Waltham, MA, USA). HEK293T cells were transfected using polyethylenimine with 500 ng of either a wildtype or mutant minigene construct and harvested 48 hours post transfection. Transfections were performed in duplicate.

### Transcript analysis *in vitro* minigene splice assay

To assess splicing total RNA was isolated using the Nucleospin RNA kit (Macherey-Nagel, Düren, Germany) following the manufacturer’s instructions. One microgram of total RNA was used as input for cDNA synthesis using iScript cDNA synthesis kit (Bio-Rad, Hercules, CA, USA) according to the manufacturer’s protocol. The region of interest was amplified from the synthesized cDNA from both wildtype and mutant minigenes using a forward primer and reverse primer located in exons 3 and 5 of the *RHO* gene. Primers amplifying *ACTB* were used as control. PCR fragments were sequence-validated by Sanger sequencing. Sequences of all primers used are listed in Supplementary table 1.

### Generation of photoreceptor precursor cells and transcript analysis

A fresh heparin blood sample was collected from the proband, and peripheral blood mononuclear cells were isolated and subsequently reprogrammed into induced pluripotent stem cells through episomal nucleofection as described previously by Koolen et al., 2022 at the Stem Cell Technology Center of the Radboudumc (Nijmegen, the Netherlands)^29^. Induced pluripotent stem cells were differentiated into PPCs by following the first 30 days of the 2D-to-3D differentiation protocol as described by Gonzalez-Cordero^30^. Cycloheximide (0.1 mg/mL Sigma-Aldrich, St. Louis, MO, USA) was added on day 30 and cells were harvested six hours post addition of cycloheximide in 350 µL RA1 buffer (Macherey-Nagel, Düren, Germany). RNA was isolated using the Nucleospin RNA kit (Machery-Nagel, Düren, Germany) following the manufacturer’s instructions.

One microgram of total RNA was used as input for cDNA synthesis using SuperScript VILO mastermix (Life Technologies, Carlsbad, CA, USA) according to the manufacturer’s protocol. Diluted cDNA was used as input for PCR (according to standard protocols) to assess the effect on *USH2A* transcripts. Sequences of all primers used are listed in Supplementary table 1.

Semi-quantitative band analysis was performed using Image Lab (Bio-Rad). Lanes were manually defined to ensure consistent alignment across samples. Bands were automatically detected and applied consistently across all gels. Background correction was performed using lane-based background subtraction.

### Nasal epithelial cell collection and targeted longread cDNA transcript analysis

Nasal epithelial cells were collected by gently swabbing the inferior turbinate epithelium using a Cytette Snap-Tip Cervical Brush (Cancer Diagnostics Inc.; Ref. SPH000, Lot 1333298). Following sampling, the brush was immediately transferred into RA1 lysis buffer supplied with the NucleoSpin RNA kit, allowing direct lysis of the epithelial material and stabilization of RNA. RNA was isolated using the Nucleospin RNA kit (Machery-Nagel, Düren, Germany) following the manufacturer’s instructions. 250 nanogram of total RNA was used as input for cDNA synthesis using SuperScript IV (Life Technologies, Carlsbad, CA, USA) and random hexamers following an adjusted protocol to enrich for long cDNA molecules. Specifically, input RNA was incubated at 60°C for 10 min to allow for the linearization of long molecules. Moreover, an extended cDNA incubation was performed at 55°C for 50 min instead of the recommended 10 min.

Following size confirmation through gel electrophoresis, PCR products were quantified using Qubit (Thermo Fisher Scientific). Targeted long-read cDNA transcript analysis was performed as described by Capasso et al., 2025^31^.

Semi-quantitative fragment analysis was performed using Image Lab (Bio-Rad). Lanes were manually assigned to ensure consistent alignment across samples. Fragments were automatically detected, applied to all gels. Background correction was performed using lane-based background subtraction.

### Estimating probability of a *de novo* single-nucleotide variant in deep intronic region of *USH2A*

The probability of a de *novo* SNV occurring in the deep-intronic region of *USH2A* could be estimated using the average genome-wide mutation rate *μ* = 1.1 · 10^−8^ per position per haploid genome^32^.

First, we calculated the size of the mutational target. Based on the canonical *USH2A* transcript ENST00000307340.8, the total genomic span is 800,558 nt and the cDNA length is 18,938 nt, resulting in an estimated total intronic length of 781,620 nt. The deep-intronic sequence was defined as the sequence located more than 100 nt from an exon boundary. After excluding 200 nt per intron across 71 introns, the deep-intronic target size was estimated as 767,420 nt. All transcript information was obtained from Ensembl (GRCh38).

Assuming independent mutation events across positions and accounting for both parental alleles, the probability of at least one *de novo* SNV was calculated using a Poisson distribution for mutation counts:

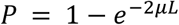

Where *μ* is the mutation rate and *L* is the deep-intronic target size.

### Zebrafish Ethics, Maintenance and Husbandry

Animal experiments were conducted in accordance with the Dutch guidelines for the care and use of laboratory animals (Wet op de Dierproeven 1996) and European regulations (Directive 86/609/EEC), as approved by the Dutch Ethics committee of the Central Committee Animal Experimentation (Centrale Commissie Dierproeven [CCD]; application number AVD10300 2022 15892). In this study we used wildtype Tupfel Longfin (TL) zebrafish, that were previously used to generate a humanized *ush2a* c.7595-2144A>G model, termed *ush2a^PE40+M^*^1 33^. In addition, we employed the previously generated knock-out *ush2a^rmc^*^1^ mutant (c.2337_2342delinsAC; p.Cys780Glnfs*32) and strain matched wildtypes^13^. Zebrafish were maintained and raised according to standard methods ^34^. Both adult and larval zebrafish were kept at a light-dark regime of 14 hours of light and 10 hours of darkness (300 lux white light). Adult zebrafish were daily fed twice with Gemma Micro 300 dry pellets (#13177, Zebcare, Nederweert, The Netherlands) at ∼5% body weight and once with artemia. Zebrafish eggs were obtained from natural spawning and reared at 28.5°C in E3 embryo medium (5 mM NaCl, 0.17 mM KCl, 0.33 mM CaCl_2_ and 0.33 mM MgSO_4_), supplemented with methylene blue.

### Immunohistochemistry, histology and quantification of fluorescent signal intensity

For the usherin staining, zebrafish larvae (5 dpf) (*ush2a^PE40+M^*^1^ and *ush2a^rmc^*^1^ (knock-out) and strain-matched wildtypes) were cryoprotected in 10% sucrose in PBS for 5 min, before embedding in OCT compound (Tissue-Tek®, #4583, Sakura, Alphen aan de Rijn, the Netherlands). Embedded samples were frozen using liquid-nitrogen-cooled isopentane and 7 µm cryosections were prepared using standard protocols.

The unfixed cryosections of zebrafish larvae were washed in PBS and permeabilized for 20 min with 0.01% Tween-20 in PBS and blocked for 30-60 min with blocking buffer (10% normal goat serum and 2% bovine serum albumin in PBS). Primary and secondary antibodies were diluted in blocking buffer. Unfixed cryosections were incubated overnight at 4°C with primary antibodies, washed (6 times for 5 min) with PBS and incubated for 1 hour at room temperature with secondary antibodies, together with DAPI (1:8000; D1306; Thermo Fisher, Waltham, MA, USA). Following a wash (6 times for 5 min) in PBS, the unfixed cryosections were post fixed with 4% paraformaldehyde for 10 min, washed (6 times for 5 min) in PBS and mounted with Prolong Gold Anti-fade (P36930; Thermo Fisher, Waltham, MA, USA). The following primary antibodies were used: rabbit anti-usherin (500 µg/mL, 1:1000; #DZ01481, Boster Bio, Pleasanton, CA, USA), and mouse anti-centrin (1:500; #04-1624; Millipore, Darmstadt, Germany). Secondary antibodies used: Alexa Fluor 568 donkey anti-rabbit (Thermo Fisher, Waltham, MA, USA, #A-11011) and Alexa Fluor 488 goat anti-mouse (Thermo Fisher, Waltham, MA, USA, #A-11029) were both used in a 1:800 dilution. All slides were imaged using a Zeiss Axio Imager fluorescence microscope equipped with an AxioCam MCR5 camera (Zeiss, Jena, Germany). Fluorescent signal intensity of anti-usherin immunoreactivity quantification was performed using Fiji version 1.53t. as described previously^35^. Means were taken and analyzed using a one-way ANOVA followed by a Tukey’s multiple comparisons test.

For rhodopsin labeling, zebrafish larvae were raised under normal light conditions (300 lux white light in a 14/10 hour day/night rhythm). At 6 dpf, larvae were sampled in the morning for exactly 100 min following light exposure. Zebrafish larvae were fixed overnight in 4% paraformaldehyde (PFA) and 100% methanol, and embedded in OCT compound (Tissue-Tek®, #4583, Sakura, Alphen aan de Rijn, The Netherlands). 7 µm sections were made, after antigen retrieval with 10mM Sodium Citrate pH8.5 at 121°C and blocking for 1h with 10% non-fat dry milk, they were stained with the primary antibody: mouse anti-rhodopsin (1:4,000, #NBP2-59690, Novus Biologicals, Centennial, CO, USA). The secondary antibody used was Alexa Fluor 488 goat anti-mouse (1:800, Thermo Fisher, Waltham, MA, USA, #A-11029) as described previously^36^. All slides were imaged using a Zeiss Axio Imager fluorescence microscope equipped with an AxioCam MCR5 camera (Zeiss, Jena, Germany). Pictures were blinded and manually counted by two individuals; averages were taken and analyzed using a non-parametric one-way ANOVA (Kruskal-Wallis) followed by a Dunn’s multiple comparison test.

## Results

### Whole genome sequencing reveals potential disease-causing variants

Individual II-2 was clinically diagnosed with a typical USH type 2 phenotype, which remained genetically unexplained after initial IDT xGen® ID Panel sequencing. The analysis revealed a pathogenic heterozygous variant in *CRB1* (NM_201253.3) c.1743_1755dup (p.(Ser586Leufs*7)), as well as two compound heterozygous pathogenic variants in *STRC* (NM_153700.2), c.2303_2313+1delGGAAGCTGCAGG (p.(?)) and c.5125A>G (p.(Thr1709Ala)) (**Figure 1**). Pathogenic variants in *STRC* are associated with mild-to-moderate autosomal recessive non-syndromic hearing loss (DFNB16) and are likely the underlying cause of hearing loss in this proband. Subsequent WGS to identify the molecular cause for RP revealed two rare compound heterozygous deep-intronic variants in *USH2A* (NM_206933.3); the previously described c.5573-834A>G and the novel c.11047+345A>G (*de novo*) variant. Variant c.5573-834A>G was previously shown to affect splicing^37^. The novel c.11047+345A>G in *USH2A* is absent from the control populations in gnomAD and is predicted to create a novel splice acceptor site near a natural existing cryptic splice donor site, leading to the potential inclusion of a 252-nt pseudoexon harboring a premature stop codon, between exons 56 and 57 of *USH2A* transcripts (p.(Val3684Thrfs*12)) (**Figure 2A**). Trio long-read-WGS confirmed the findings from short-read WGS, Sanger-based segregation analysis and *STRC*-specific testing and did not reveal additional candidate variants, in particular no potentially pathogenic variant *in trans* to the known *CRB1* variant.

**Figure 1:**
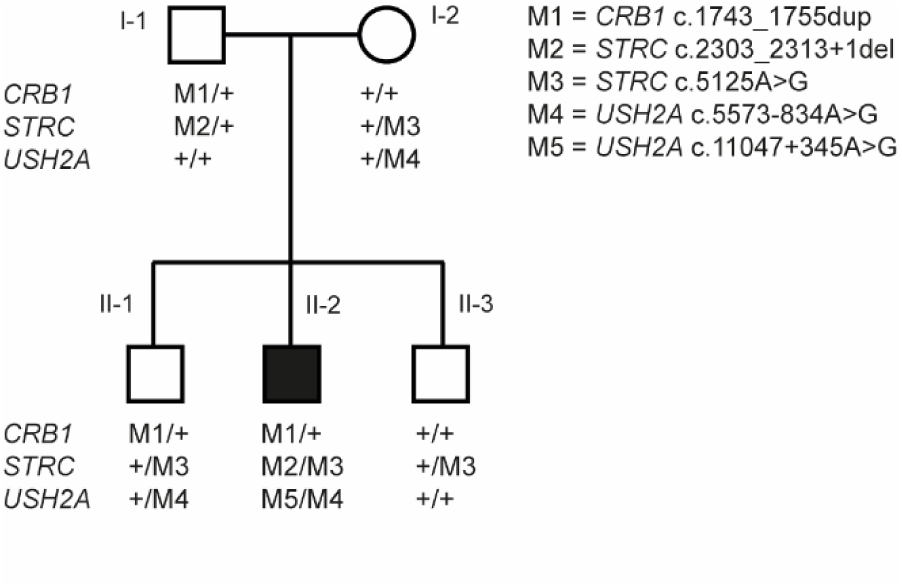
Pedigree showing the segregation of variants in *CRB1*, *STRC*, and *USH2A*. Squares represent males and circles represent females, the filled symbol indicates the affected individual (II-2). For each family member, the tested variants are shown per gene, with “+” indicating the wildtype allele. M5 is absent from both parents.

**Figure 2:**
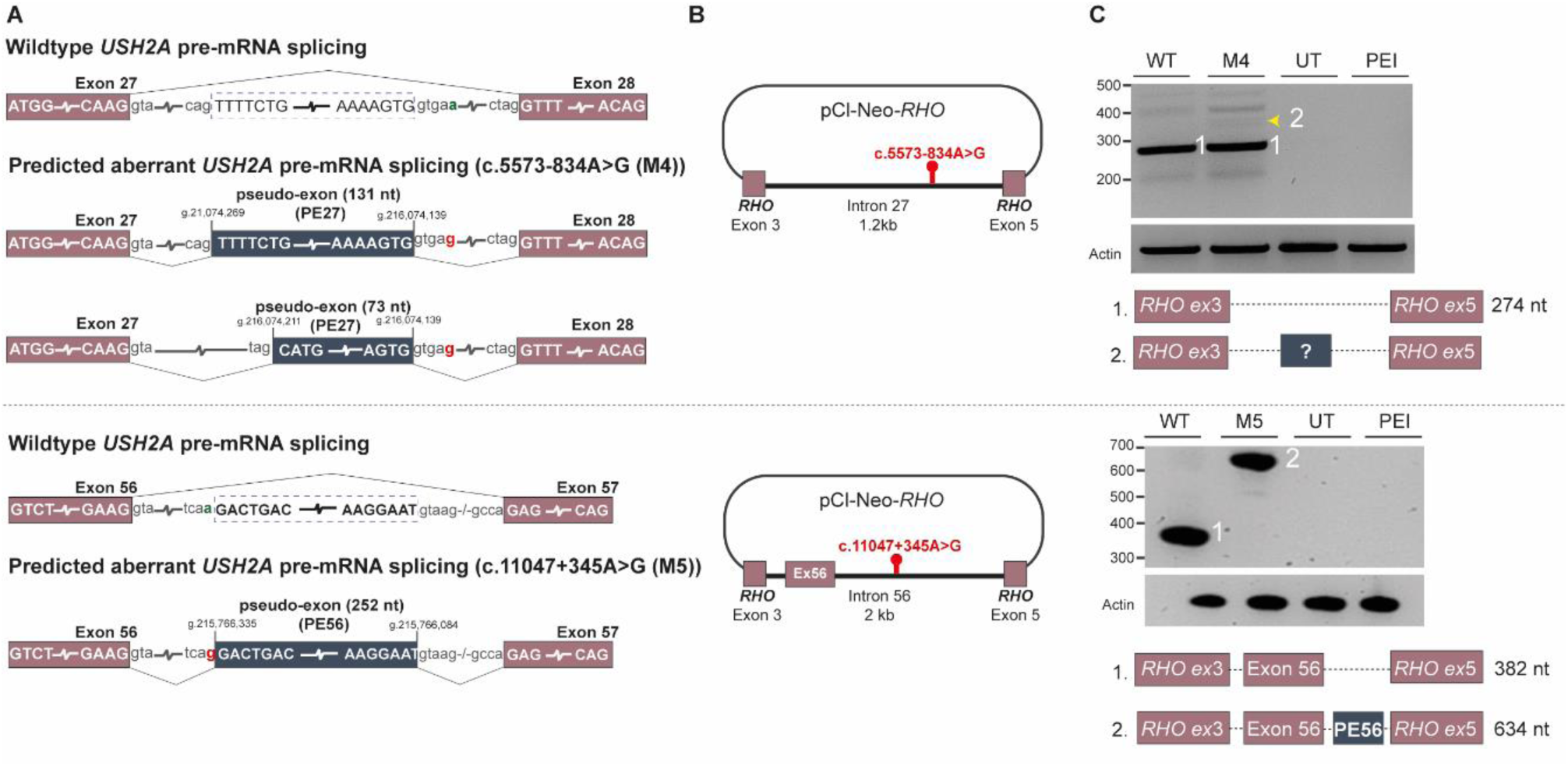
Minigene splice assay analysis of the *USH2A* c.5573-834A>G (M4) and c.11047+345A>G (M5) variants. **(A)** Predicted splicing effects of the deep-intronic variants c.5573-834A>G (M4) and c.11047+345A>G (M5), showing activation of cryptic splice sites and potential inclusion of pseudoexons between the flanking exons. **(B)** Schematic overview of the minigene constructs containing *USH2A* intron 27 (c.5573-834A>G) or exon 56 + intron 56 (c.11047+345A>G) fragments cloned into the pCI-Neo-*RHO* exon 3-exon 5 backbone. **(C)** RT-PCR analysis of transcripts generated from the *USH2A* minigene constructs following transfection in HEK293T cells. Presence of the c.5573-834A>G variant only results in a very faint additional fragment (yellow arrowhead), and does not result in clearly detectable pseudoexon inclusion. The presence of the c.11047+345A>G variant results in a larger fragment corresponding to the predicted pseudoexon inclusion. Amplification of the housekeeping gene *Actin* is used as a loading control. WT: wildtype, UT: untransfected control, PE: pseudoexon, PEI: polyethylenimine, Ex: exon, nt: nucleotide.

### Minigene analysis reveals variant-specific splicing effects

We performed minigene splice assays in HEK293T cells using our previously published pCI-Neo-*RHO* exon 3-exon 5 vector to assess the potential effect of the c.11047+345A>G variant on splicing, as well as to recapitulate the findings of the previously described c.5573-834A>G variant^28^. For each variant, the flanking genomic region was PCR-amplified from proband genomic DNA, and the resulting fragments were then recombined into the expression vector using Gateway cloning to produce the wildtype and mutant constructs (**Figure 2B**).

This minigene splice assay for the *USH2A* c.5573-834A>G variant did not confirm the previously described partial inclusion of two pseudoexons (73 and 131 nt), as no obvious evidence of aberrant pre-mRNA splicing was observed and only a very faint, non-informative additional fragment was present (**Figure 2C**). Additionally, using gel electrophoresis followed by Sanger sequencing, we confirmed the inclusion of a 252-nt pseudoexon resulting from the *de novo* variant c.11047+345A>G, leading to premature termination of usherin translation, p.Val3684Thrfs*12 (Fig. 2C).

### Aberrant splicing of c.5573-834A>G confirmed in patient-derived photoreceptor cells

As our results did not correspond with the previously reported results of aberrant pre-mRNA splicing induced by the c.5573-834A>G variant in HEK293T cells, we next assessed the effect of both variants on *USH2A* pre-mRNA splicing in a more physiologically relevant cellular model. Using patient-derived photoreceptor precursor cells (PPCs) that contain both *USH2A* variants within the genetic context of the affected individual, we performed transcript analyses using primers designed for the flanking exons of both *USH2A* deep-intronic variants.

For variant c.5573-834A>G, transcript analysis in PPCs treated with NMD inhibitor cycloheximide showed evidence for the presence of both wildtype as well as aberrantly spliced *USH2A* transcripts (**Figure 3A**). Semi-quantitative fragment analysis indicates that pseudoexon inclusion occurs in a minority of transcripts derived from the c.5573-834A>G allele. Approximately 18.8% of transcripts include the aberrant splice product, while approximately 81.2% retains the wildtype configuration. Sanger sequence analysis of the aberrant spliced transcripts confirmed the inclusion of a 131-nt pseudoexon. The predicted 73nt pseudoexon was not observed. While for variant c.11047+345A>G the HEK293T-based splice assay showed evidence for the inclusion of a pseudoexon, direct mRNA analysis from patient-derived PPCs did not provide evidence for the presence of pseudoexon-containing *USH2A* transcripts resulting from this variant (**Figure 3A**).

**Figure 3:**
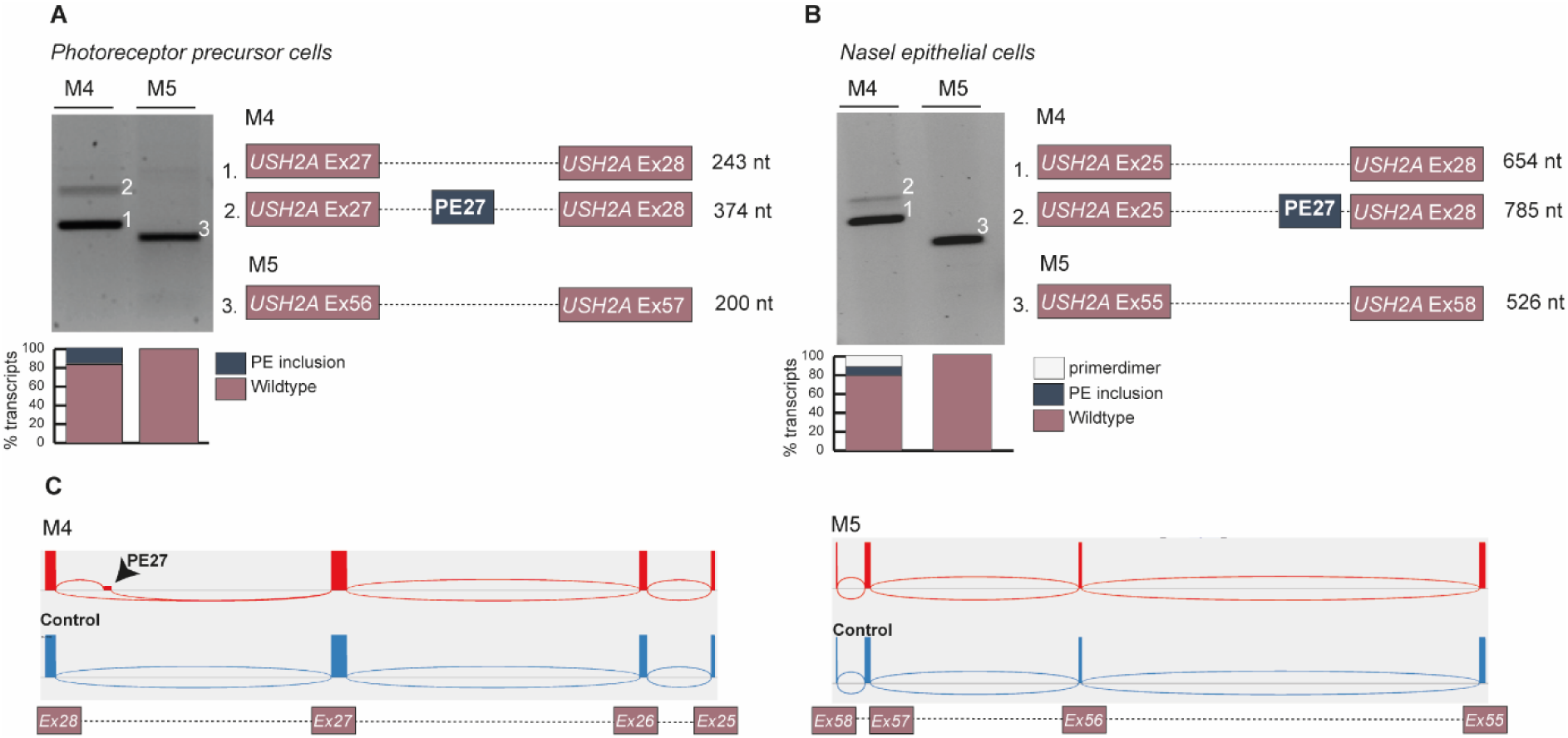
RT-PCR analysis of pseudoexon inclusion caused by the *USH2A* c.5573-834A>G (M4) and c.11047+345A>G (M5) variants in patient-derived photoreceptor precursor cells and nasal epithelial cells. **(A)** RT-PCR analysis of *USH2A* transcripts spanning exons 27-28 and exons 56-57 in patient-derived photoreceptor precursor cells carrying the M4 and M5 variants. Both wildtype fragments and larger fragments corresponding to pseudoexon inclusion are detected for M4. For M5 only wildtype fragments are detected. Corresponding exon structures and expected product sizes are shown below the gel. **(B)** RT-PCR analysis of *USH2A* transcripts spanning exons 25-28 and exons 55-58 in patient-derived nasal epithelial cells carrying the M4 and M5 variants. As for panel A, both wildtype fragments and larger fragments representing pseudoexon inclusion are detected for M4. For M5 only wildtype fragments are detected **(C)** Targeted long-read PacBio RNA sequencing performed on nasal epithelial RNA confirmed inclusion of the 131-nt pseudoexon associated with the M4 variant. The Sashimi plot supports clear splice-junctions for the aberrant exon-pseudoexon-exon splicing of M4, confirming the RT-PCR findings.

### Nasal epithelial cells recapitulate splice alterations observed in photoreceptor precursor cells

We extended the analysis by investigating the effect of both deep-intronic variants on *USH2A* pre-mRNA splicing in patient-derived nasal RNA collected. As was done in the previous experiment, we performed a direct analysis of nasal mRNA using primers that were designed for the exons flanking both *USH2A* deep-intronic variants.

The transcript analysis from patient-derived nasal epithelial cells, showed evidence for the inclusion of a 131-nt pseudoexon in approximately 6.7% of transcripts related to variant c.5573-834A>G, and approximately 79.7% of wildtype transcripts (13.6% primer-dimer was observed) (**Figure 3B**), based on semi-quantitative band analysis. For variant c.11047+345A>G no evidence of aberrantly spliced *USH2A* pre-mRNA was obtained (**Figure 3B**). To complement these findings, we performed targeted long-read PacBio RNA sequencing on the nasal RNA. The Sashimi plot generated from this dataset confirmed inclusion of the 131-nt pseudoexon in a subset of transcripts carrying variant c.5573-834A>G, whereas no aberrant *USH2A* splicing events were detected resulting from variant c.11047+345A>G (**Figure 3C**). These long-read data further support the observations obtained from both nasal epithelial cells and PPCs.

Collectively, results obtained from minigene splice assays differ clearly from the data obtained from the studies in patient-derived nasal epithelia and PPCs. In addition, the observed effect of the previously published c.5573-834A>G on aberrant *USH2A* pre-mRNA splicing is a partial defect, further casting caution on the pathogenicity of both variants.

### Partial PE40 inclusion in zebrafish leads to reduced usherin signal

In order to gain insights in the level of residual wildtype *USH2A* transcripts that are enough to produce sufficient usherin protein for normal retinal function, we adopted the humanized *ush2a^PE40+M^*^1^ knock-in zebrafish line. A human *USH2A* intron 40 cassette (termed PE40+M1) was previously inserted in the zebrafish *ush2a* gene between exons 40 and 41^33^. This cassette was optimized for recognition by the zebrafish splicing machinery and includes the deep-intronic c.7595-2144A>G variant. Quantitative transcript analyses in retinae of this zebrafish line revealed that 89.1 ± 3.3% of all *ush2a* transcripts contained the 152 nt human PE40 sequence, resulting in a frameshift and a premature termination of usherin translation. In addition, 10.9 ± 3.3% of all *ush2a* transcripts are still fully wildtype. To determine the consequences at the level of usherin protein expression, we performed an immunohistochemical analysis on wildtype, *ush2a^PE40+M^*^1^ and *ush2a^rmc^*^1^ (knock-out) larvae (5 dpf) using a previously validated anti-zebrafish usherin antibody. Usherin is located at the periciliary membrane of wildtype zebrafish photoreceptor cells in close vicinity of the connecting cilium and basal body marker centrin. As expected, no usherin was detected in the retina of *ush2a^rmc^*^1^ larvae (**Figure 4A**). In the retina of *ush2a^PE40+M^*^1^ larvae, usherin was still detected, although the intensity of the anti-usherin signal appeared lower than that in wildtype larvae. Quantification of the fluorescence intensity signals using an automated Fiji script confirmed that usherin immunoreactivity was significantly reduced in *ush2a^PE40+M^*^1^ larvae (36.51 ± 7.59) as compared to wildtype controls (61.11 ± 20.90) (P ≤ 0.0001, one-way ANOVA followed by Tukey’s multiple comparison test) (**Figure 4B**). Comparison of the usherin immunoreactivity in the *ush2a^PE40+M^*^1^ retina with that in the retina of *ush2a^rmc^*^1^ mutants (13.08 ± 4.03) revealed a significantly higher signal (P ≤ 0.0001, one-way ANOVA followed by Tukey’s multiple comparison test).

**Figure 4:**
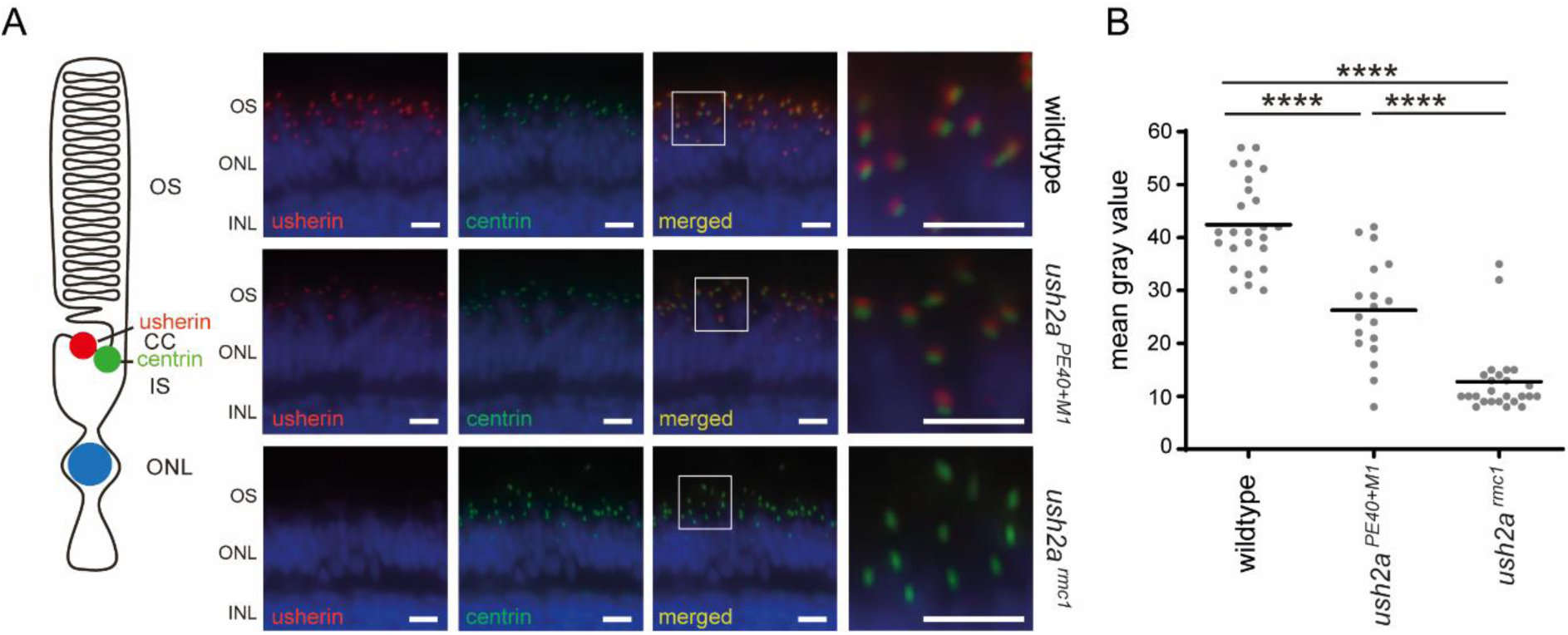
Visualization of usherin in retinal sections of wildtype, *ush2a^PE40+M^*^1^ and *ush2a^rmc^*^1^ larvae. **(A)** Retinal cryosections of wildtype, *ush2a^PE40+M^*^1^ and *ush2a^rmc^*^1^ larvae (5 days post fertilization) labeled with antibodies directed against usherin (red) and centrin (green). Nuclei are counterstained with DAPI (blue). Scale bar: 10 μm. OS: outer segment, CC: connecting cilium, IS: inner segment, ONL: outer nuclear layer. **(B)** Quantification of anti-usherin signal intensity at the photoreceptor periciliary membrane. Individual data points represent the mean fluorescence intensity of anti-usherin signals within a single middle section of one larval zebrafish eye. Horizonal bars depict the mean signal intensity within a genotype (n = 25-28 eyes per genotype). Data were analyzed using one-way ANOVA followed by Tukey’s multiple comparison test. **** P ≤ 0.0001.

### Minimal residual wildtype transcript is sufficient to maintain near normal photoreceptor function in zebrafish

Previous research has shown that one of the most important roles of usherin in photoreceptor cells is supporting rhodopsin trafficking from the photoreceptor inner-to-outer segments^38^. To assess whether the observed level of residual wildtype usherin expression in *ush2a^PE40+M^*^1^ larvae is sufficient for normal photoreceptor function, we evaluated the distribution of rhodopsin in all three zebrafish models. Cryosections were made from wildtype (n=23), *ush2a^PE40+M^*^1^ (n=17) and *ush2a^rmc^*^1^ (n=22) larvae (6 dpf). While aberrant distribution of rhodopsin was prominently observed in photoreceptors of *ush2a^rmc^*^1^ larvae (11.31 ± 6.16 photoreceptors with aberrant rhodopsin localization/cryosection), rhodopsin was mostly correctly localized to the rod photoreceptor outer segments *ush2a^PE40+M^*^1^ larval eyes (6.37 ± 4.46 photoreceptors with aberrant rhodopsin localization/cryosection), with no significant difference and at levels comparable to those in wildtype larval eyes (4.37 ± 4.69 photoreceptors with aberrant rhodopsin localization/cryosection) (**Figure 5**). This suggests that, in zebrafish, only 10.9 ± 3.3% of wildtype *ush2a* transcript is still sufficient to produce enough usherin protein for near normal rhodopsin trafficking. Translating these findings to the human situation, this would imply that significant care should be taken in the pathogenicity assessment of (deep-intronic) genetic variants that only have a partial effect on aberrant pre-mRNA *USH2A* splicing.

**Figure 5:**
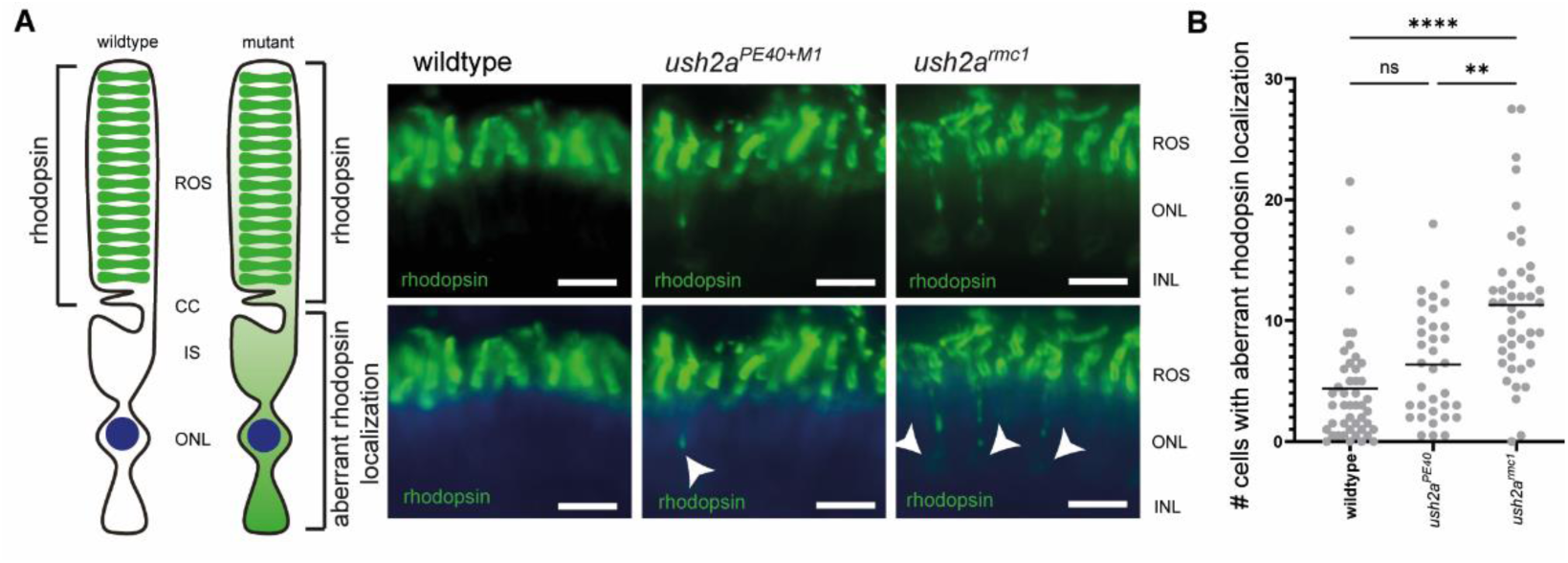
Visualization of rhodopsin localization in retinal sections of wildtype, *ush2a^PE40+M^*^1^ and *ush2a^rmc^*^1^ larvae. **(A)** Retinal cryosections of wildtype, *ush2a^PE40+M^*^1^ and *ush2a^rmc^*^1^ larvae (6 days post fertilization) labeled with antibodies directed against rhodopsin (green). In wildtype larvae, rhodopsin localizes correctly to the rod outer segment, whereas *ush2a^rmc^*^1^ larvae display prominent aberrant rhodopsin distribution (arrowheads). Rhodopsin localization in *ush2a^PE40+M^*^1^ larvae is largely comparable to wildtype. Nuclei are counterstained with DAPI (blue). Scale bar: 10 μm. ROS: Rod outer segment, CC: connecting cilium, IS: inner segment, ONL: outer nuclear layer. **(B)** Quantification of photoreceptors with aberrant rhodopsin localization in wildtype (n=23), *ush2a^PE40+M^*^1^ (n=17) and *ush2a^rmc^*^1^ (n=22) larvae. Individual data points represent the number of affected photoreceptors within one larval zebrafish eye. Horizontal bars depict the mean number of affected cells within a genotype. Data were analyzed using a non-parametric one-way ANOVA (Kruskal-Wallis) followed by Dunn’s multiple comparison test. ** and **** indicate significant differences between groups, ns indicates non-significance.

## Discussion

Multiple splice-affecting variants in *USH2A* have been identified as the underlying cause of *USH2A*-associated disease. While certain variants have been shown to fully abolish correct pre-mRNA splicing, others result in the inclusion of a pseudoexon in only part of the transcripts, resulting in both aberrantly and wildtype spliced *USH2A* transcripts. For the latter variants, the impact on protein function remains unclear, and these variants are typically classified as VUS following the ACMG classification guidelines, hindering a conclusive genetic diagnosis in the current USH case.

Here, we studied two deep-intronic variants, c.5573-834A>G and c.11047+345A>G, using an *in vitro* minigene splice assay, patient-derived PPCs and nasal epithelial cells. Through the use of an *in vitro* minigene splice assay we observed the inclusion of a 252-nt pseudoexon for the *de novo* variant c.11047+345A>G. *USH2A* variant c.5573-834A>G was previously tested through an *in vitro* minigene splicing assay and was shown to result in aberrant pre-mRNA splicing, while also retaining correctly spliced *USH2A* transcripts^37^. The aberrantly spliced mRNA transcripts were shown to include either a pseudoexon fragment of 73 nt or a fragment of 131 nt, both resulting from the activation of a cryptic donor site. In contrast to these previously published results, through our *in vitro* minigene assay we only observed a very faint, non-informative fragment. Although *in vitro* minigene assays are widely used, relatively easy to conduct and cost-effective as a first-line method for the evaluation of splice-altering variants, it is an artificial system in which the outcomes might depend on the design of the assay (flanking region length, inclusion of flanking exons, backbone choice), execution (amount of DNA used in transfection, time of harvesting) and host cells used. This variability in design and execution probably underlies the difference between the previously reported results for variant c.5573-834A>G and the results in this study. In addition, the *in vitro* minigene assay was performed in HEK293T cells that lack retina-specific splice factors, potentially influencing the outcome of the analyses. These limitations potentially contribute to the different outcomes between the minigene splice assay and the patient-derived PPCs.

To combine a more physiologically relevant context with the entire genetic context of the patient, we performed transcript analysis of mRNA isolated from both patient-derived PPCs as well as nasal epithelial cells. While variant c.5573-834A>G was shown to result in incomplete inclusion of a pseudoexon in both models used, only normally spliced mRNA was observed for the *de novo* variant c.11047+345A>G in these systems. Because aberrantly spliced transcripts that introduce a premature termination codon are typically degraded by nonsense-mediated decay, aberrant products are potentially underrepresented in systems where NMD is active. In PPCs, cycloheximide was added to inhibit NMD. In nasal epithelial cells, where NMD cannot be blocked, the pseudoexon-containing fragment appears to be produced at similarly low levels as in cycloheximide-treated PPCs. Taken together, both patient-derived model systems confirmed the presence of pseudoexon inclusion resulting from the c.5573-834A>G variant in only a small fraction of *USH2A* transcripts.

To enable a better classification of splice-altering variants with a partial effect on pre-mRNA gene splicing, we used a previously generated humanized zebrafish model, which was initially designed to assess antisense oligonucleotide-based splice modulation therapies for the pathogenic c.7595-2144A>G *USH2A* variant. In USH patients, variant c.7595-2144A>G leads to complete incorporation of PE40 in the *USH2A* transcripts^10^. In contrast, in the humanized zebrafish model PE40 is incorporated in ∼85% of *ush2a* transcripts ^33^. This model is therefore suitable for examining the impact of residual wildtype *ush2a* transcripts on usherin expression and function and provides additional knowledge guiding human *USH2A* variant interpretation and classification.

While we did observe reduced levels of usherin at the photoreceptor periciliary membrane of *ush2a^PE40+M^*^1^ larvae compared to strain- and age-matched wildtype controls, the level of usherin expression was still significantly higher in *ush2a^PE40+M^*^1^ larvae as compared to the level of usherin expression in the complete knockout zebrafish *ush2a^rmc^*^1^ model. These data suggest that a minimal amount of residual wildtype transcripts is sufficient to result in the expression of detectable levels of usherin in the retina of *ush2a^PE40+M^*^1^ zebrafish. We emphasize the translational value of the zebrafish model for interpreting *USH2A* variants. Dulla et al. (2021) showed that the in-frame skipping of *ush2a* exon 13 in roughly 20% of transcripts was sufficient for a complete rescue of zebrafish retinal function in their assays^12^. In the same study, treatment of patient-derived PPCs and mouse models with antisense oligonucleotides (including QR-421a/ultevursen) resulted in a dose-dependent skipping of *USH2A* exon13 skipping. Based on these studies, patients have been treated with a dose of QR-421a/ultevursen predicted to result in *USH2A* exon13 skipping levels of ∼20-30% of transcripts. Early clinical data from the phase 1b/2a Stellar trial (#NCT03780257) showed early signs of efficacy following QR-421a/ultevursen treatment, and this approach is currently being evaluated in the phase 2b LUNA trial (#NCT06627179). These preclinical and early clinical findings support the hypothesis that even a partial restoration of *USH2A* transcript/protein levels can be functionally and clinically meaningful.

Building further on these data, we propose a dose-sensitive model for interpreting *USH2A* splice-altering variants. We suggest that pathogenicity is only expected when residual wildtype transcript levels drop below a functional threshold (∼15-20% of the wildtype *USH2A* transcript levels in healthy individuals. Because semi-quantitative fragment analysis did not indicate a severe loss of wildtype transcript, as the wildtype transcript represents the clear majority (∼80%) in both PPC and nasal-cell assays, we classify c.5573-834A>G as a VUS rather than a (likely) pathogenic variant. Additionally, we consider the *de novo* variant c.11047+345A>G a (likely) benign variant as only correctly spliced *USH2A* transcripts were observed in patient-derived material. To further strengthen this classification, we estimated the probability of encountering any *de novo* SNV within the deep-intronic region of *USH2A* using the average genome-wide mutation rate (*μ* = 1.1×10⁻⁸ per base per haploid genome)^32^. Based on the calculated deep-intronic target size (767,420 nt), the probability of at least one *de novo* SNV occurring in this region was approximately 1 in 60 births (0.0167 or ≈1,67%). Although this estimate does not account for sequence context or predicted pathogenicity, it provides an order-of-magnitude indication of how likely such a *de novo* event is expected to be. In combination with the absence of aberrant splicing in patient material, this suggests that the identified *de novo* variant is unlikely to have functional consequences and therefore most consistent with a (likely) benign interpretation. Finally, we recommend obtaining quantitative transcript measurements, segregation data, and additional functional assays before assigning a genetic diagnosis for *USH2A*-related retinal disease. Biallelic variants in *STRC* typically result in recessively inherited moderate hearing loss (DFNB16), and could therefore attribute to the hearing loss observed in our patient. However, care should be taken as the recorded hearing loss is more severe than what is typically seen in DFNB16-affected individuals. Therefore the presence of an additional, yet undetected, disease-modifying variant could be expected. Because neither short- nor long-read WGS identified a potentially pathogenic variant *in trans* to the pathogenic *CRB1* variant c.1743_1755dup (p.(Ser586Leufs*7)), it most likely represents carriership that is unrelated to the probands retinal phenotype. Furthermore, these data warrant that the results obtained from an *in vitro* pre-mRNA splicing assays for *USH2A* should be interpreted with care as the mere presence of an alternatively spliced transcript does not necessarily make the variant causative for *USH2A*-associated retinal disease.

## Acknowledgements

The authors want to thank the proband and his family members for his participation in this study. The authors would like to thank the Radboud Genome Technology Center for infrastructural and computational support and the Radboudumc Stem Cell Technology Center for reprogramming and characterizing the patient line. The authors would also like to thank Ellen Blokland and Marlie Jacobs-Camps for DNA sample preparation and administration. The authors would like to thank Antoon van der Horst and Jeroen Boerrigter for technical assistance and acknowledge the Radboud University Zebrafish Facility for excellent zebrafish husbandry. We would also like to thank Gavin Arno for his expertise and valuable advice in optimizing the nasal epithelial cell collection procedure.

## Data availability statement

Data are available upon reasonable request. All other genome sequencing data are subject to controlled access because they may compromise the privacy of research participants. These data may become available upon a data transfer agreement approved by the local ethics committee and can be obtained after contacting the corresponding author upon request.

## Supplementary tables

**Supplementary table 1.**
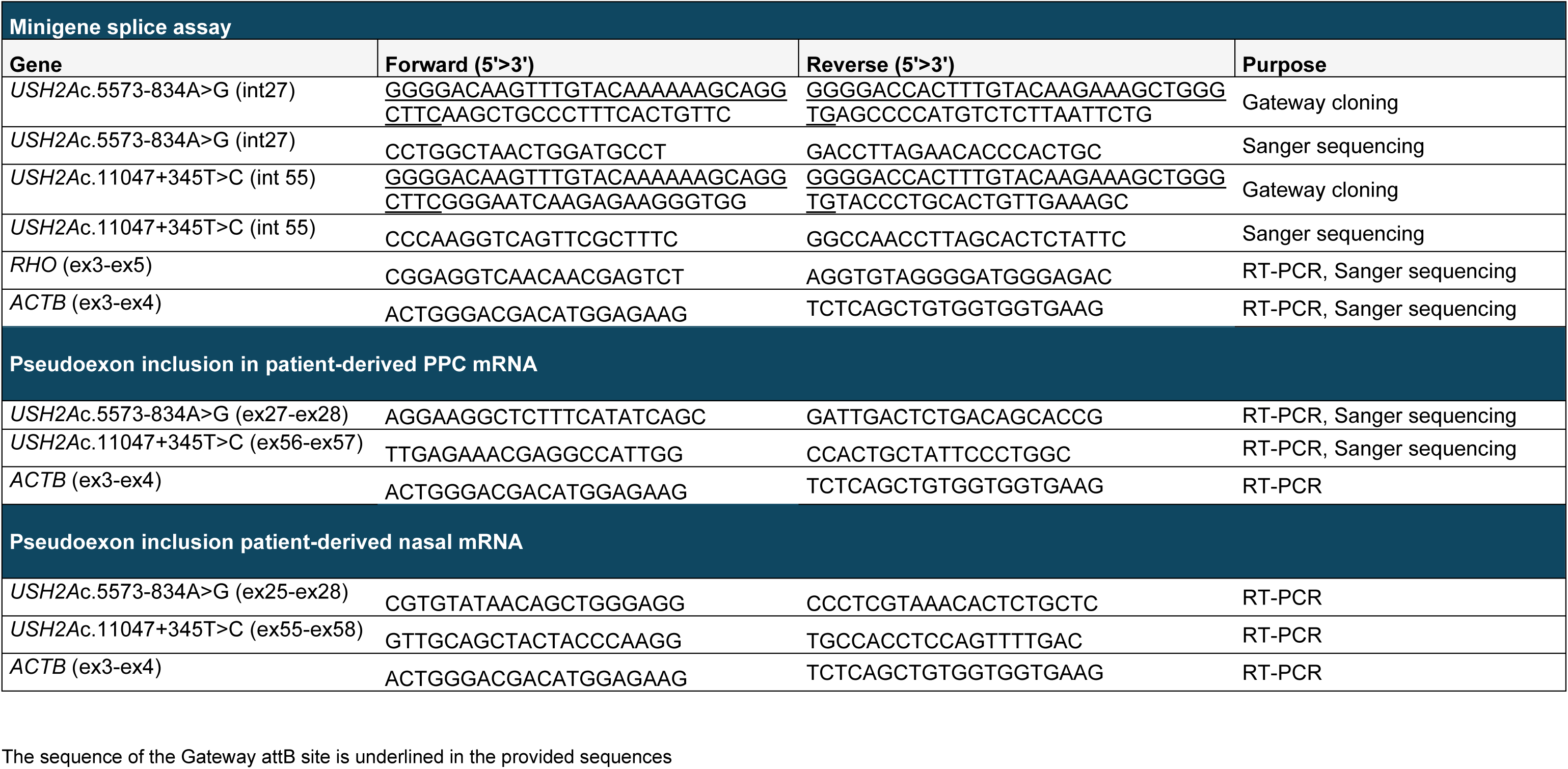
Overview of oligonucleotide sequences used in this study.

